# Gas vesicle-blood interactions enhance ultrasound imaging contrast

**DOI:** 10.1101/2023.07.24.550434

**Authors:** Bill Ling, Jeong Hoon Ko, Benjamin Stordy, Yuwei Zhang, Tighe F. Didden, Dina Malounda, Margaret B. Swift, Warren C.W. Chan, Mikhail G. Shapiro

## Abstract

Gas vesicles (GVs) are genetically encoded, air-filled protein nanostructures of broad interest for biomedical research and clinical applications, acting as imaging and therapeutic agents for ultrasound, magnetic resonance, and optical techniques. However, the biomedical applications of GVs as a systemically injectable nanomaterial have been hindered by a lack of understanding of GVs’ interactions with blood components, which can significantly impact *in vivo* performance. Here, we investigate the dynamics of GVs in the bloodstream using a combination of ultrasound and optical imaging, surface functionalization, flow cytometry, and mass spectrometry. We find that erythrocytes and serum proteins bind to GVs and shape their acoustic response, circulation time, and immunogenicity. We show that by modifying the GV surface, we can alter these interactions and thereby modify GVs’ *in vivo* performance. These results provide critical insights for the development of GVs as agents for nanomedicine.

## INTRODUCTION

Nanomaterials are becoming increasingly important for biomedical applications such as drug delivery, medical imaging, and diagnostics^1^. In these contexts, nanoparticle behavior is significantly impacted by cells and proteins encountered within the bloodstream. Serum proteins rapidly adsorb to nanoparticle surfaces, forming a protein corona that alters their physicochemical properties and recognition by the body^2–4^. The corona’s composition can predict factors such as pharmacokinetics, biodistribution, toxicity, and cellular uptake^5–7^. Modification strategies often involve covering the particle surface with hydrophilic polymers such as polyethylene glycol (PEG) and other ligands^8^. Additionally, some nanomaterials bind to erythrocytes (RBCs), affecting imaging contrast^9^, biodistribution^10^, and circulation time^11^.

Gas vesicles (GVs) are an emerging nanomaterial with great potential as agents for imaging and therapy^12^. These air-filled protein nanostructures are naturally produced by certain aquatic microbes for buoyancy regulation^13^. GVs comprise a 2-nm thick protein shell that excludes liquid water but permits the dynamic exchange of gas, forming a thermodynamically stable pocket of air with nanoscale dimensions^13^. Acoustic waves are strongly scattered at this air-water interface, enabling GVs to produce robust ultrasound contrast when injected into the body^14–16^ or expressed in engineered cells^17,18^. Furthermore, they are resilient to repeated insonation^14^, easily tailored to target molecular markers^19–21^ or respond to biological functions^22,23^, and have growing applications in therapeutic ultrasound^24,25^, optical imaging^26,27^, and magnetic resonance imaging^28,29^. To effectively incorporate these capabilities into an injectable agent, a deeper understanding of *in vivo* GV behavior is needed.

In this study, we investigate GV interactions with RBCs and serum proteins, develop surface functionalization techniques to modulate these interactions, and evaluate the downstream effects on acoustic response, circulation time, and immunogenicity. We characterize GVs’ protein corona and identify molecular pathways governing their *in vivo* fate. This analysis offers valuable insights for the ongoing development and optimization of injectable nanoparticle and GV-based agents.

## RESULTS

### Gas vesicles adsorb to red blood cells

We began by studying the behavior of GVs after intravenous (IV) administration. We visualized circulating GVs with ultrafast power Doppler ultrasound imaging, leveraging their ability to enhance blood flow contrast^15^. Targeting a single coronal plane in the mouse brain, we acquired images at a 15.625 MHz center frequency and 0.25 Hz frame rate (**Fig. 1A**). After a 5 min baseline, we injected 100 µL of 5.7 nM GVs purified from *Anabaena flos-aquae*^30^ and monitored the ensuing changes in hemodynamic signal. In healthy BALB/c mice, contrast reached an initial peak within 1 min, followed by a larger peak 3.5 min later, then returned to baseline over approximately 30 min as GVs were cleared by the liver^23^ (**Fig. 1B**). Intensities at the first peak were consistent across trials but varied significantly at the second peak (**Fig. 1C**).

**Figure 1.**
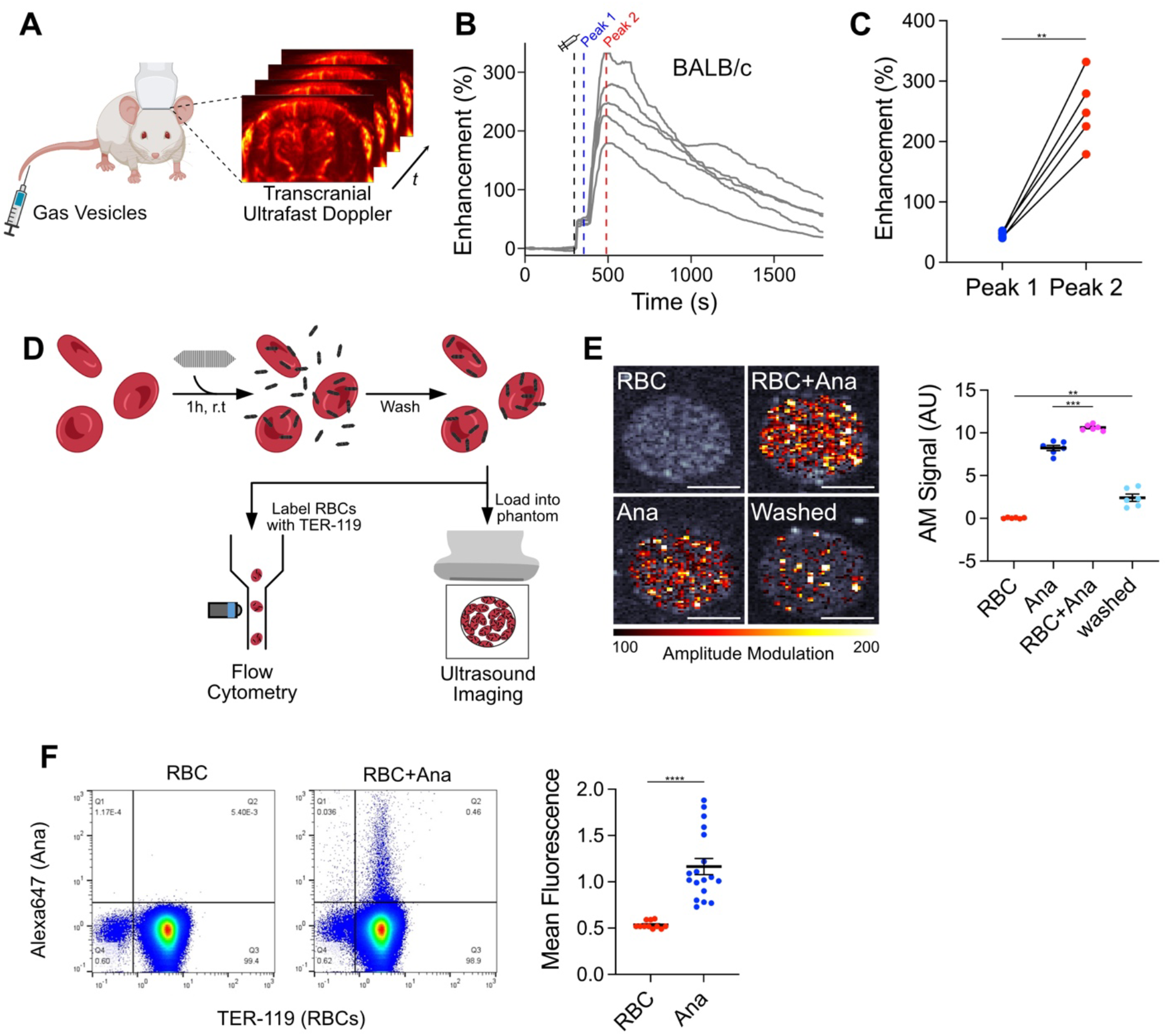
GV adsorption to RBCs contributes to a second peak of hemodynamic ultrasound contrast. **(A)** Diagram of *in vivo* imaging setup. Intravascular dynamics of IV injected GVs were visualized by transcranial ultrafast power Doppler imaging of the brain. **(B)** Time courses of Doppler signal enhancement in immunocompetent BALB/c mice following injection of 100 µL 5.7 nM GVs. N = 5. Dashed gray line, time of injection (300 s); dashed blue line, peak 1 (350 s); dashed red line, peak 2 (480 s). **(C)** Signal enhancement at the indicated peaks in time courses from panel (B). Points from the same trial are connected. N = 5. Paired t-test, (**, p < 0.01). **(D)** Diagram of RBC binding assay. Ultrasound imaging: RBCs were incubated with GVs modified to produce non-linear signal, washed by centrifugation, and loaded into an agarose phantom for nonlinear AM imaging. Flow cytometry: RBCs were incubated with fluorescently-labeled GVs, washed by centrifugation, stained with anti-TER-119, and analyzed by flow cytometry. **(E)** Acoustic detection of adsorbed GVs. Left: Representative ultrasound images of RBCs mixed with GVs. AM signal is overlaid on a B-mode image to show sample outlines. Scale bars, 1 mm. Right: Mean AM signal intensity within each well. N = 6. Error bars, ± SEM. Welch’s t-test, (**, p < 0.01; ***, p < 0.001). **(F)** Flow cytometric detection of GVs adsorbed to RBCs. Left: Representative scatter plots of washed RBCs, gated for single cells. Gating strategy is shown in Fig. S3. Right: Mean fluorescence of TER-119+ cell population. RBC, N = 11; RBC+Ana, N = 18. Error bars, ± SEM. Welch’s t-test, (****, p < 0.0001).

We next investigated the cause of this dual-peak phenomenon. We hypothesized that the first peak was due to dispersion of free-floating GVs throughout the bloodstream, as this process is expected to occur within 15 seconds of injection^31^, while the second peak could arise from an increase in acoustic backscatter due to GV clustering^14^. We observed similar contrast enhancement dynamics in immunocompetent BALB/c (**Fig. 1B**) and immunocompromised NSG mice (**Fig. S1**), and therefore suspected an antibody-independent mechanism such as adsorption to RBCs, as previously seen with nanobubbles^9^. To evaluate this concept, we calculated the theoretical scattering cross-section^32^ of RBC-GV complexes. Modeled as uniform spheres with volume-weighted physical properties, scattering cross-section increased with the number of adsorbed GVs and was greater than that of dispersed particles (**Fig. S2**).

To quantify adsorption, we exposed purified mouse RBCs to GVs that were modified to produce nonlinear ultrasound contrast^21^. The RBCs were maintained at 10% of *in vivo* levels to facilitate uniform mixing, while the GVs were at concentrations approximating *in vivo* conditions following vascular dispersion. After 1 h, we loaded the samples into an imaging phantom and detected GVs specifically with an amplitude modulation (AM) pulse sequence^33^ (**Fig. 1D**). Signal intensities for RBC and GV controls were 0.04 AU and 8.24 AU, respectively, and increased to 10.62 AU after mixing. After centrifugation to remove unbound GVs, 23% (2.41 AU) of this signal was retained (**Fig. 1E**).

We validated these results by incubating RBCs with GVs labeled with a fluorescent dye. After 1 h, we washed the cells thoroughly to remove loosely bound GVs and analyzed them by flow cytometry (**Fig. 1D, S3**). Mean fluorescence of the population doubled from 0.53 to 1.17, with 0.5% of RBCs showing significant binding (**Fig. 1F**). Taken together, our data suggest that increased acoustic backscatter from GV adsorption to RBCs contributes to the delayed wave of hemodynamic contrast. This mechanism may operate in concert with others such as serum-induced aggregation, which we examine below.

### PEG-coated GVs do not interact with RBCs

To minimize RBC adsorption, we engineered GVs coated with methoxypolyethylene glycol (mPEG), a widely-used polymer for nanoparticle passivation^8^. We functionalized the GV surface with alkyne groups (**Fig. S4**) and attached 10 kDa mPEG-azides through a copper-catalyzed azide-alkyne cycloaddition (CuAAC) (**Fig. 2A**). We will refer to unmodified GVs as Ana, and to functionalized GVs as Ana-PEG. Consistent with the addition of a PEG layer, dynamic light scattering (DLS) showed an increase in hydrodynamic diameter from 240 nm to 370 nm (**Fig. 2B**), while zeta potential neutralized from −56 mV to −5 mV (**Fig. 2C**). We next performed pressurized absorbance spectroscopy, which tracks optical density under increasing hydrostatic pressure to determine the threshold at which GVs collapse, providing a convenient measure of structural integrity^14,30^. Ana and Ana-PEG collapsed at 600 kPa and 450 kPa, respectively, suggesting that attachment of mPEG mildly destabilized the GV shell, but that most of its strength was intact (**Fig. 2D**). Incubation with mPEG or CuAAC reagents separately had no effect (**Fig. S5**), while direct functionalization with NHS-PEG severely compromised shell stability and failed to shield surface charge (**Fig. S6**). B-mode ultrasound contrast from both GV types was equivalent (**Fig. 2E**).

**Figure 2.**
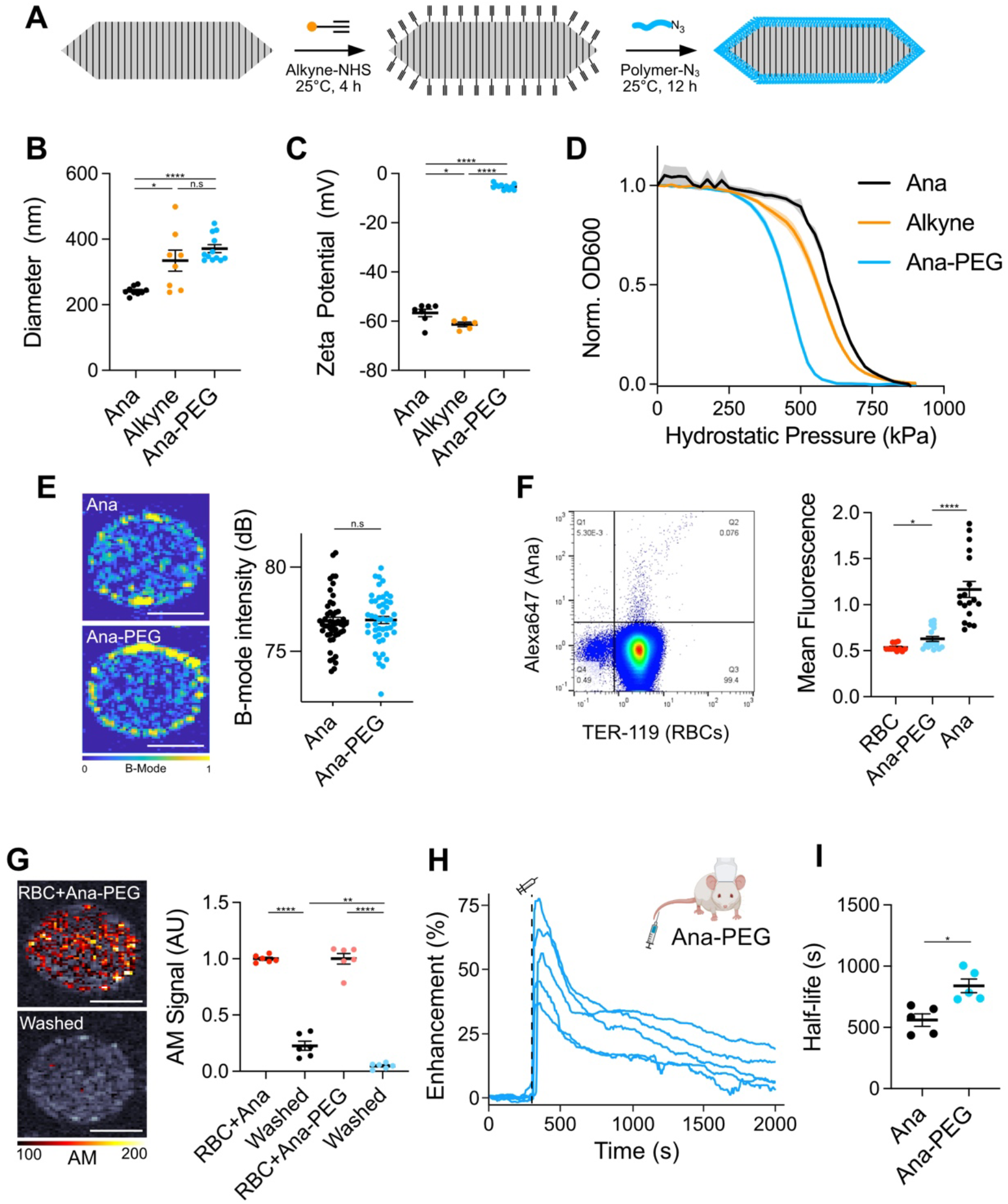
Surface passivation reduces RBC binding and extends circulation time. **(A)** Reaction scheme for GV functionalization. Alkynes were conjugated to lysines on the GV surface, and polymers were attached through a CuAAC reaction. **(B)** DLS measurements of hydrodynamic diameter. N = 8−12. Error bars, ± SEM. Welch’s t-test, (*, p < 0.05; ****, p < 0.0001; n.s, p ≥ 0.05). **(C)** Zeta potential measurements of engineered GVs. N = 5−11. Error bars, ± SEM. Welch’s t-test, (*, p < 0.05; ****, p < 0.0001). **(D)** Normalized optical density at 600 nm as a function of hydrostatic pressure. N = 4. Thick lines, mean; shaded areas, ± SEM. **(E)** Left: Representative B-mode images of Ana and Ana-PEG embedded in an agarose phantom. Right: Mean B-mode signal intensities within each well. N = 48. Error bars, ± SEM. Welch’s t-test, (n.s, p ≥ 0.05). **(F)** Flow cytometric detection of fluorescently-labeled Ana-PEG adsorbed to RBCs. Left: Representative dot plot of washed RBCs, gated for single cells. RBCs are stained with anti-TER-119. Right: Mean fluorescence of TER-119+ cell population. N = 18. Ana and RBC-only controls from Fig. 1F are shown as a reference. Error bars, ± SEM. Welch’s t-test, (*, p < 0.05; ****, p < 0.0001). **(G)** Acoustic detection of Ana-PEG modified to produce nonlinear contrast. Left: Representative ultrasound images of RBCs mixed with Ana-PEG. AM signal is overlaid on a B-mode image. Right: AM signal intensities, normalized to their respective washed samples. N = 6. Normalized data from Fig. 1E are shown for comparison. Error bars, ± SEM. Welch’s t-test, (**, p < 0.01; ****, p < 0.0001). **(H)** Time courses of ultrafast power Doppler signal enhancement following injection of Ana-PEG into BALB/c mice. N = 5. Dashed line, time of injection (300 s). **(I)** Half-life of GV-induced signal enhancement calculated by fitting time courses in Fig. 1B (Ana) and Fig. 2H (Ana-PEG) to an exponential decay function. Error bars, ± SEM. Welch’s t-test, (*, p < 0.05).

Next, we evaluated the effectiveness of this coating at reducing RBC adsorption. As before, we mixed purified mouse RBCs with fluorescently-labeled Ana-PEG for 1 h, removed loosely bound GVs by centrifugation, and analyzed the cells by flow cytometry (**Fig. 1D**). Less than 0.1% of cells exhibited significant binding, with mean fluorescence of the population only increasing to 0.63, compared to 1.17 for Ana (**Fig. 2F**). Likewise, less than 5% of ultrasound signal was retained after washing away unbound Ana-PEG, compared to 23% with Ana (**Fig. 2G**).

Having confirmed the reduced adsorption of Ana-PEG to RBCs, we assessed their *in vivo* behavior. Following IV injection of 100 µL 5.7 nM Ana-PEG into healthy BALB/c mice, hemodynamic contrast reached a maximum within 1 min before returning to baseline monotonically (**Fig. 2H**). The timing and magnitude of enhancement at this peak was consistent with the first peak observed after Ana injection (**Fig. 1B**), supporting our hypothesis that this initial peak resulted from vascular distribution. Unlike in the Ana time course, however, a second peak of contrast enhancement was not observed. Fitting these time courses to an exponential decay model, we found that PEGylation increased apparent circulation half-life from 560 s to 840 s (**Fig. 2I**).

### Serum protein-mediated aggregation

GV aggregation is an alternative mechanism to increasing acoustic backscatter^14^ which cannot be excluded by the results presented thus far. Due to their highly charged surfaces (**Fig. 2C**), GV aggregation is unlikely to occur spontaneously^34^ and would most likely be facilitated by a component within serum. To test this idea, we incubated Ana and Ana-PEG in 80% serum from naïve BALB/c, NSG, and outbred non-Swiss mice for 1 h at 37°C and measured flotation, a reliable indicator of clustering^14^, by comparing optical densities at the surface and in the subnatant (**Fig. 3A**). We included NSG mice due to their lack of antibodies, while the genetic heterogeneity of outbred mice increases the likelihood of forming native antibodies^35^ against GVs. Prior to incubation, the optical density ratio was 1.1 for both GV types and increased only slightly in BALB/c and NSG serum (without statistical significance) (**Fig. 3B**). Upon exposure to outbred serum, Ana formed a distinct buoyant layer (ratio 1.9), while Ana-PEG showed a less pronounced increase (ratio 1.3). Pre-incubation transmission electron microscopy (TEM) images contained only discrete particles (**Fig. 3C**); exposure to outbred serum caused Ana to assemble into multi-GV bundles, whereas Ana-PEG formed occasional small clusters but remained mostly dispersed.

**Figure 3.**
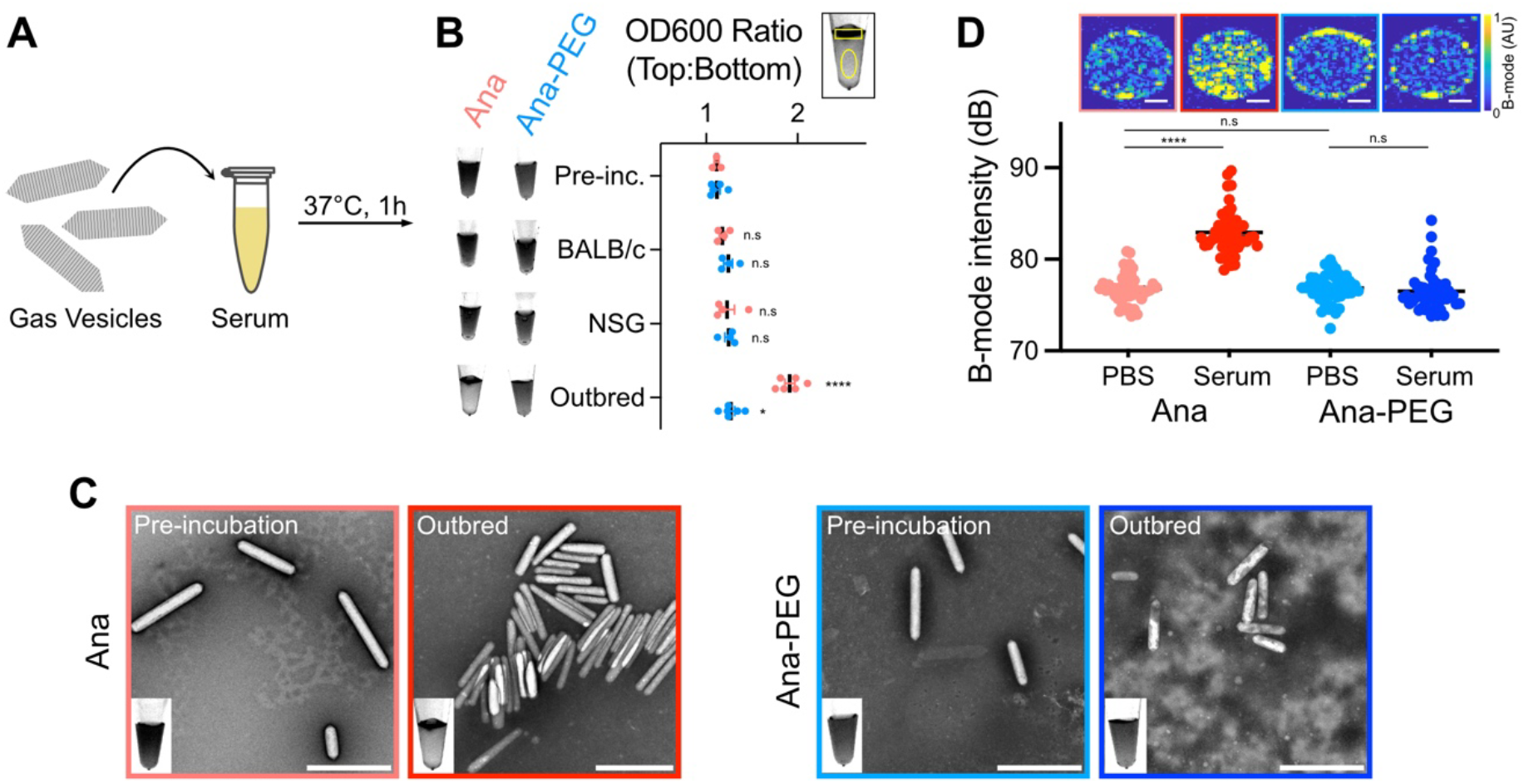
Serum exposure can cause GVs to aggregate. **(A)** Diagram of serum incubation assay. GVs (0.23 nM) were incubated in 80% mouse serum for 1 h at 37°C. **(B)** Optical detection of GV flotation. Left: Representative trans-illumination images of serum-incubated GVs. Right: Ratio of optical densities in manually drawn ROIs at the surface and in the subnatant of each sample. Representative ROIs are shown above the plot. Error bars, ± SEM. Welch’s t-test, compared to pre-incubation samples (*, p < 0.05; ****, p < 0.0001; n.s, p ≥ 0.05). **(C)** Representative TEM images of GVs before and after exposure to serum from outbred mice. Inset shows trans-illumination image of corresponding sample. Scale bars, 500 µm. **(D)** Ultrasound imaging of GVs following incubation in PBS or outbred mouse serum. Top: Representative B-mode images. Scale bars, 1 mm. Bottom: Mean signal intensity within each well. N = 48. Error bars, ± SEM. Welch’s t-test, (****, p < 0.0001; n.s, p ≥ 0.05).

To compare acoustic responses, we embedded GVs treated with either PBS or outbred serum into an imaging phantom and acquired ultrasound images using a B-mode pulse sequence (**Fig. 3D**). Signal intensity from Ana increased by 6.13 dB in outbred serum relative to PBS, while contrast from Ana-PEG remained relatively unchanged. In BALB/c serum, signal from Ana increased by 2.26 dB, consistent with a lower degree of aggregation compared to outbred serum (**Fig. S7**).

### Immunogenicity and effect of antibodies in vivo

We next investigated the impact of elicited antibodies on GV dynamics *in vivo* by administering multiple GV injections to the same animals. We injected BALB/c mice with an initial dose of 100 µL 5.7 nM GVs and did so again 1 or 4 weeks later (**Fig. 4A**). Ana injections resulted in peak enhancements of approximately 280% at all three timepoints (**Fig. 4B−C**), with apparent circulation half-life decreasing from 560 s to 290 s and 410 s at weeks 1 and 4, respectively (**Fig. 4D**). Repeated Ana injections were well-tolerated, with no health anomalies observed by veterinary assessment.

**Figure 4.**
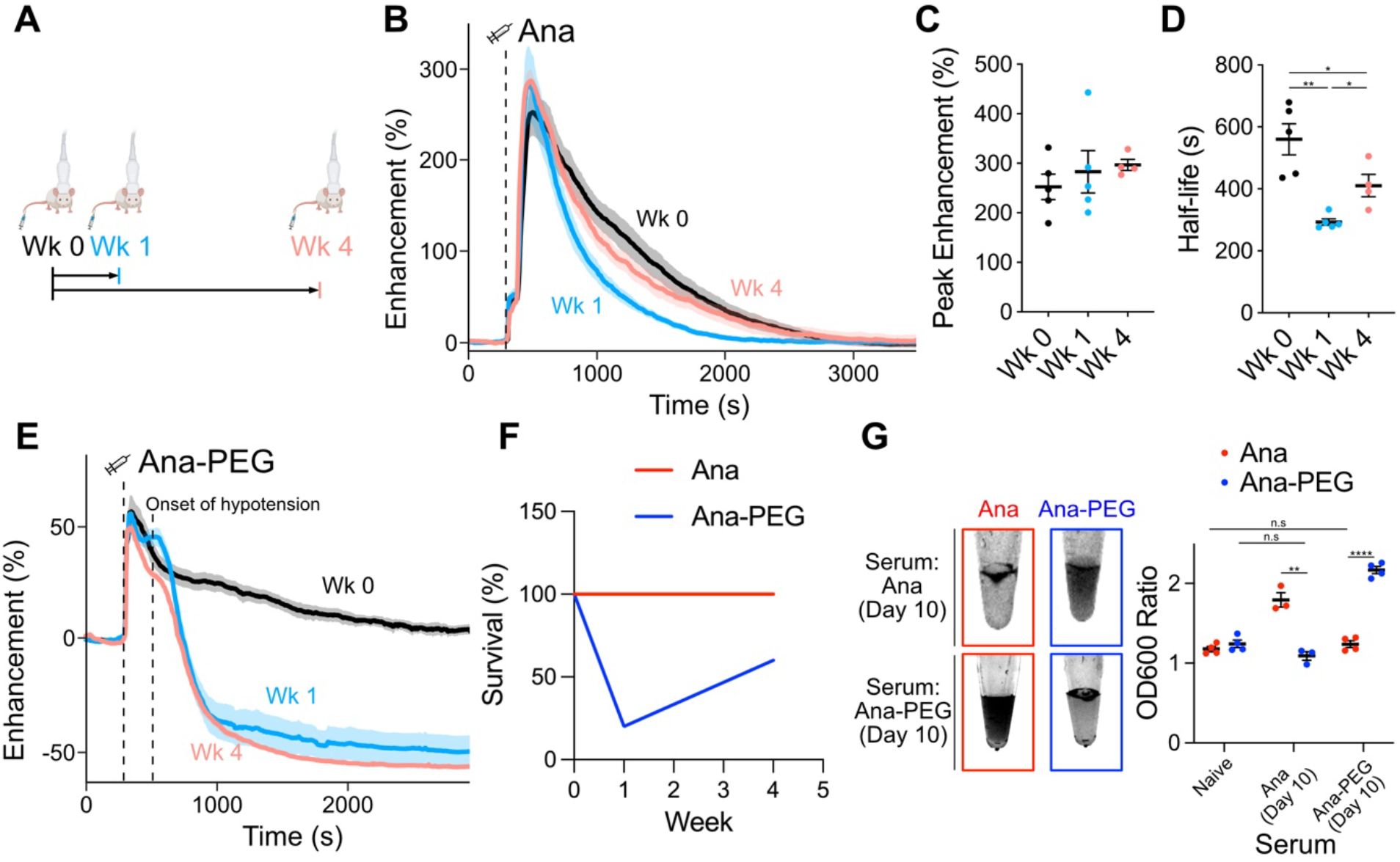
Antibody reactions to GVs. **(A)** Timeline of GV injections. Immunocompetent mice were injected with two doses of 100 µL 5.7 nM Ana or Ana-PEG, separated by either 1 or 4 weeks. Ultrafast Doppler imaging was performed at each injection. **(B)** Time courses of hemodynamic signal enhancement following administration of Ana. Thick lines, mean; shaded area, ± SEM; dashed line, time of injection (300 s). N = 4-5. **(C)** Peak enhancement of time courses in panel B. Error bars, ± SEM. **(D)** Circulation half-life calculated by fitting time courses in panel B to an exponential decay function. Error bars, ± SEM. Welch’s t-test, (*, p < 0.05; **, p < 0.01). **(E)** Time courses of hemodynamic signal enhancement following administration of Ana-PEG. Severe hypotension occurred within 5 min of injection, resulting in a sharp drop in hemodynamic signal. Thick lines, mean; shaded area, ± SEM; dashed lines, time of injection (300 s) or onset of hypotension (500 s). N = 4-5. **(F)** Survival rate following GV administration. All mice dosed with Ana recovered after both injections. Several mice dosed with Ana-PEG did not recover after the second injection. N = 5 at each time point. **(G)** GV aggregation in the presence of anti-GV antibodies. Left: Representative trans-illumination images of GVs incubated in serum prepared from mice 10 days post-immunization. Right: Ratio of optical density at the surface relative to the subnatant. Values from incubation in naïve BALB/c serum (Fig. 3B) are included for comparison. N = 3-4. Error bars, ± SEM. Welch’s t-test, (**, p < 0.01; ****, p < 0.0001; n.s, p ≥ 0.05).

Ana-PEG produced peak enhancements of approximately 50% at all three timepoints (**Fig. 4E, S8**). However, acute hypotension occurred unexpectedly several minutes after the second dose of Ana-PEG, resulting in a sharp reduction in hemodynamic contrast (**Fig. 4E**). Only 20% of mice recovered from hypotension at week 1 and 60% at week 4 (**Fig. 4F**). Based on similar responses to other PEGylated materials^36,37^, we hypothesized that this reaction is triggered by anti-PEG antibodies.

To further investigate antibody interactions, we exposed GVs to serum prepared from mice 10 days after the initial GV injection (**Fig. 4G**). In the presence of Ana-PEG antiserum, Ana-PEG aggregated and formed a buoyant layer within 30 min, while Ana remained in suspension. Conversely, Ana rapidly aggregated upon exposure to Ana antiserum, while Ana-PEG remained in suspension. This pattern is consistent with a clear distinction in antibody selectivity and minimal cross-reactivity for the different GV surfaces. Substituting mPEG with a 16 kDa zwitterionic polymer, which is being explored as a less immunogenic PEG alternative^38,39^, did not alleviate coating-induced anaphylaxis (**Fig. S9**).

Taken together, our data suggest that RBC adsorption is the primary contributor to GV-enhanced hemodynamic contrast, whereas serum components mediate particle clearance and elicit immune responses. This conclusion is supported by the comparable Doppler signal dynamics observed in antibody-deficient NSG (**Fig. S1**), naïve BALB/c (**Fig. 1B**), and GV-exposed BALB/c mice (**Fig. 4B**). Furthermore, peak enhancement was not impacted by the presence of serum factors capable of causing significant GV aggregation (**Fig. 4C, G**). Instead, these factors triggered an acceleration in GV clearance or, in the case of Ana-PEG, an infusion reaction (**Fig. 4D, F**).

### Protein corona of GVs is dominated by immune response proteins

To identify the serum components influencing GV behavior, we characterized the protein coronas associated with Ana and Ana-PEG. We incubated both GV types in serum from outbred mice for 1 h at 37 °C, as it offers a more diverse representation of serum components and enhances generalizability for translational applications^40^. After removing unbound proteins by centrifugation, we digested bound proteins with trypsin and quantified peptides by liquid chromatography tandem mass spectrometry (LC-MS/MS) (**Fig. 5A**, see SI for full dataset). Levels of detected proteins showed moderate correlation between serum-exposed GV types (R = 0.77) but less so with the background serum (Ana R = 0.58, Ana-PEG R = 0.45) (**Fig. S10**), indicating that Ana and Ana-PEG selectively enrich for similar proteins through a process that cannot be explained by serum concentration alone. We detected comparable amounts of the GV structural protein GvpC across all samples, indicating similar GV loading, as well as minor quantities of GvpV, GvpN, and other cyanobacterial proteins (**Fig. S11**).

**Figure 5.**
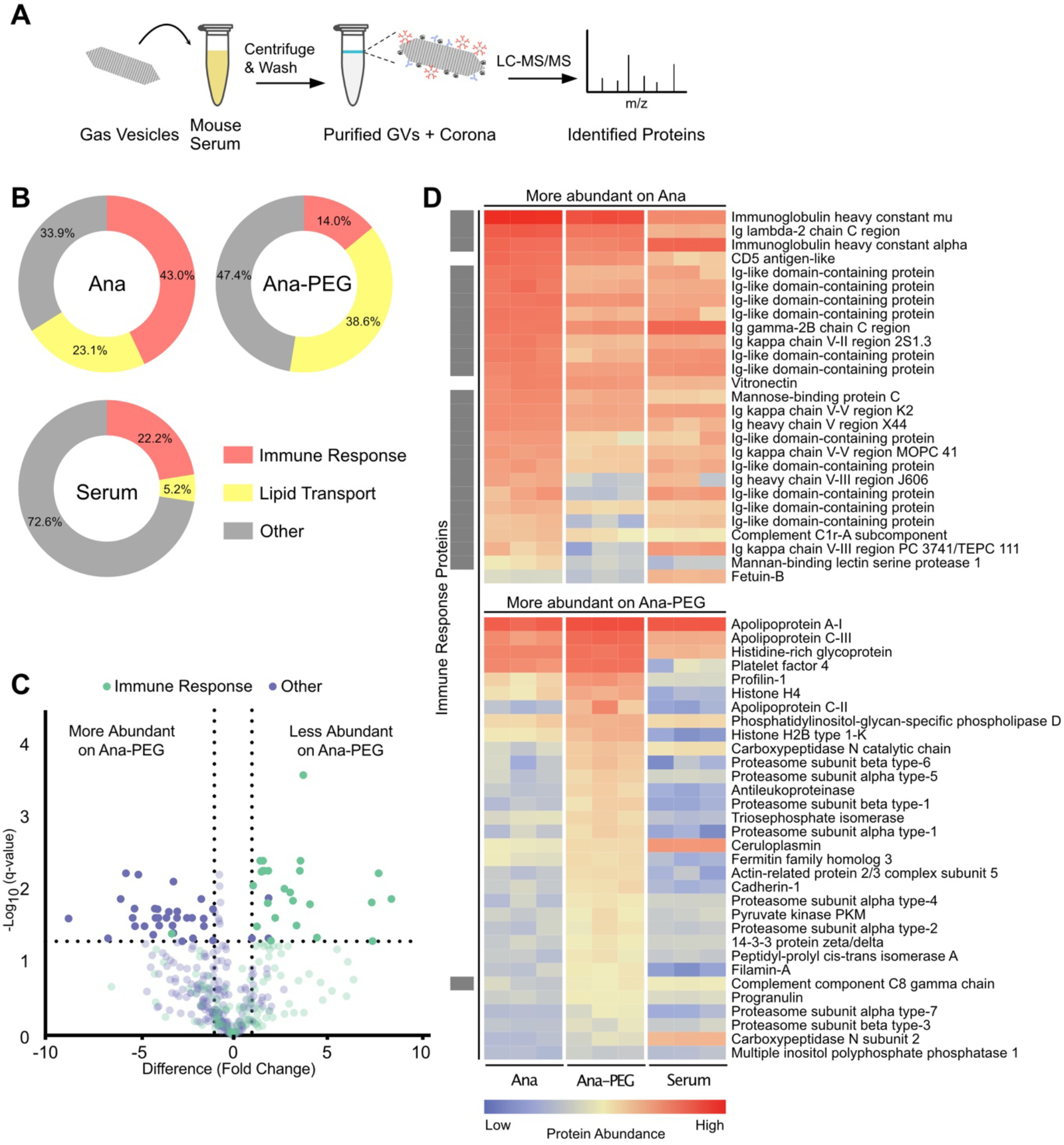
Characterization of the GV protein corona. **(A)** Schematic of corona analysis protocol. GVs were incubated in outbred mouse serum for 1 h at 37°C, purified by centrifugation, and processed for LC-MS/MS. **(B)** Donut charts of relative abundances of immune response and lipid transport proteins, as classified by gene ontology. **(C)** Volcano plot of protein abundance on Ana compared to Ana-PEG. Abundances were compared by multiple unpaired t-test analysis using the false discovery rate method of Benjamini and Hochberg. Proteins with a false discovery rate of 5% and log_2_ fold-change greater than 1 were deemed differentially abundant. Proteins that were not differentially abundant are shown translucently. **(D)** Heat map of differentially abundant proteins on Ana compared to Ana-PEG, shown with a mouse serum control. Immune response proteins are indicated by a gray box on the left.

Immune response proteins—part of an immunoglobulin complex or involved in complement activation—were highly abundant in the Ana corona, constituting 43.0% of detected proteins, compared to 14.0% of the Ana-PEG corona and 22.2% of serum (**Fig. 5B**). Of the proteins significantly more enriched on Ana than Ana-PEG, 24 of 27 were associated with immune response, including immunoglobulin A, immunoglobulin G, immunoglobulin kappa, mannose-binding protein C, and complement C1r (**Fig. 5C−D**). Similarly, 13 of the top 25 proteins enriched on Ana relative to serum were in this group (**Fig. S12**). In contrast, only one immune response protein was among the 32 proteins more abundant on Ana-PEG than Ana. Consistent with its role as an early-response antibody^41^, immunoglobulin M (IgM) was the most prevalent member of this group, accounting for 15% of the Ana corona (1^st^ overall) and 4% of the Ana-PEG corona (8^th^ overall). Given its enrichment and multivalency, IgM is likely responsible for GV aggregation in naïve serum.

Lipid transport proteins were enriched on both GV types, comprising 23.1% and 38.6% of the Ana and Ana-PEG coronas, respectively, compared to 5.2% of serum (**Fig. 5B**). Apolipoprotein E (ApoE) and apolipoprotein C-I (ApoC-I) were the most prominent. On Ana, they ranked 2^nd^ and 4^th^ in overall abundance with 130-fold and 50-fold enrichment relative to serum, respectively (**Fig. S12**). On Ana-PEG, they were 1^st^ and 2^nd^ with 200-fold and 90-fold enrichment, respectively. Ana-PEG also enriched several other proteins that are typically found in low concentration in serum, such as apolipoprotein C-II, apolipoprotein C-III, platelet factor 4, and profilin-1 (**Fig. 5D, S12**). Neither GV type appreciably adsorbed albumin despite its high concentration in serum.

## DISCUSSION

Our results demonstrate that blood components significantly influence GV behavior in the blood stream, highlighting opportunities for optimizing GV-based diagnostic and therapeutic agents. Injected GVs adsorb to the surface of RBCs, resulting in a considerable enhancement of hemodynamic contrast. Additionally, GVs acquire a serum protein corona that is rich in apolipoproteins and immunoglobulins. These corona proteins can facilitate rapid clearance and amplify the immune response upon repeated exposure. Surface passivation with mPEG reduces RBC and protein adsorption, providing a modest extension of circulation time at the expense of diminished blood flow contrast.

GV-based diagnostic agents could benefit from strategies to modify the protein corona, which can mask elements required for molecular detection and response. Potential techniques include genetic functionalization of the GV surface^21^, ligand conjugation to serum-equilibrated GVs^42^, fusion of peptides to recruit specific serum proteins^43^, and adsorption of an artificial corona^44^. These strategies could also enable *in vitro* diagnostic applications, such as clustering-based detection in which GVs selectively enriched with specific proteins are combined with the corresponding antibodies, allowing for rapid optical and acoustic measurements via flotation and enhanced ultrasound backscatter^14,45^, respectively. Furthermore, the GV corona can aid in proteomics by reducing the dynamic range of protein concentrations in biological fluids, facilitating detection of low-abundance components^46^. GVs are advantageous for these applications due to their easy buoyancy-based isolation and use of structural proteins as internal concentration standards.

To maximize the translational utility of GVs, immunogenic components should be identified and eliminated. GVs do not appear to be immunotoxic, as repeated injections of Ana were well-tolerated. However, antibodies did form against the GVs, leading to accelerated clearance. Notably, several residual cyanobacterial proteins remained after GV purification. Future work could study responses to urea-treated GVs lacking these proteins^30^, identify problematic epitopes by analyzing peptides displayed on antigen-presenting cells after lysosomal processing of the GV^47^, and redesign production and purification processes to address these challenges.

Enhancing GV binding to RBCs could potentially alleviate immunogenicity concerns by inducing peripheral tolerance^48,49^, while also extending GV circulation time^50,51^ and improving contrast in functional ultrasound imaging^15^. Approaches include covalent linkage to engineered RBCs^50^ or incorporation of RBC affinity ligands to enhance binding in situ^51^. Future work should also examine the impact of GV adsorption on RBC structure and function, including morphology, longevity, and gas exchange.

In conclusion, our study provides several key insights into GV interactions with blood components, uncovering mechanisms underlying their recognition by the body and the balance between circulation half-life and contrast enhancement. By understanding the impact of these interactions on performance and safety, we move closer to optimizing GVs as injectable imaging agents and realizing the full potential of this promising technology.

## Supporting information

Mass Spectrometry Data

## ACKNOWLEDGMENTS

The authors wish to thank Prof. Bob Grubbs, Prof. Mark E. Davis, and Dr. Di Wu for helpful discussion; Dr. Hojin Kim for help with the aqueous SEC polymer characterization; Justin Lee for assistance with flow cytometry; Dr. Mona Shahgoli for assistance with small molecule mass spectrometry; the Caltech Cryo-EM Center for assistance with TEM; the Caltech CCE Multiuser Mass Spectrometry Lab for instrumentation to characterize synthesized small molecules; Dr. Craig Simpson and Dr. Leanne Wynbenga-Groot, The Hospital for Sick Children, Toronto, Canada for assistance with mass spectrometry for proteomic analysis. This research was supported by the National Institutes of Health (R01-EB018975 to M.G.S.) and the Rosen Bioengineering Center Pilot Grant. W.C.W.C. acknowledges the Canadian Institute of Health Research Grants FDN159932 and MOP-1301431, NMIN Network 2019-T3-01 and Canadian Research Chairs Program Grant 950-223824. J.H.K was supported by the Kavli Nanoscience Institute Prize Postdoctoral Fellowship at the California Institute of Technology. B.L. was supported by the NIH/NRSA Pre-Doctoral Training Grant (T32GM07616) and the Caltech Center for Environmental and Microbial Interactions. B.S. thanks the Doctoral

Completion Award. M.G.S. is a Howard Hughes Medical Institute Investigator.

## AUTHOR CONTRIBUTIONS

.L, J.H.K., and M.G.S. conceptualized the research. B.L performed the *in vivo* imaging experiments with assistance from M.B.S. J.H.K. designed polymer synthesis and gas vesicle functionalization reactions with assistance from T.F.D. J.H.K and B.L. characterized the functionalized gas vesicles. B.L. performed the erythrocyte modeling and incubation experiments. B.S. and Y.Z. performed LC−MS/MS experiments and analyzed the data. D.M. prepared gas vesicles for experiments. All authors contributed to editing and revising the manuscript. M.G.S. and W.C.W.C. supervised the research.

## DATA AND MATERIALS AVAILABILITY

All gas vesicles, plasmids, data, and code are available from the authors upon reasonable request.

