## Supplementary Information for "Gas vesicle-blood interactions enhance ultrasound imaging contrast"

### Methods

#### Chemicals

All chemicals were purchased from Sigma-Aldrich and used without further purification unless otherwise noted.

#### GV preparation

Native gas vesicles were isolated from *Anabaena flos-aquae* as previously described[1] and stored in 1x phosphate-buffered saline (PBS). Concentrations were measured by optical density (OD) at 500 nm with a NanoDrop ND-1000 (Thermo Scientific). The following relations were used for stoichiometry calculations: OD 1 GV = 114 pM GV = 4.41 uM GvpA. GVs producing nonlinear contrast were prepared by removing GvpC as previously described[2]. Fluorescently labeled GVs were prepared by adding 90 μM Alexa Fluor 647 N-hydroxysuccinimide ester (Invitrogen, prepared as 10 mM stock solution in DMSO) to a suspension of OD 30 GVs (5000 dye per GV). After rotating gently at room temperature for 2 h, excess dye was removed by two rounds of overnight dialysis through a regenerated cellulose membrane (12-14 kDa MWCO, Repligen) followed by two rounds of centrifugation.

#### Polymer coating

GV-alkynes were prepared by adding 26.46 mM propargyl-N-hydroxysuccinimide ester (250 mM stock solution in DMSO) to a suspension of OD 50 GVs (120 alkyne/gvpA). After rocking gently for 4 h at room temperature, the GVs were purified by 4 rounds of centrifugation (600xg, 2h).

For a typical CuAAC reaction, 55.5 mg m-PEG-azide (10 kDa, BroadPharm), 110 μL DMSO, 12 μL PBS, and 54 μL aminoguanidine hydrochloride[3] (11.11 mg/mL in PBS) were combined in a 2 mL microcentrifuge tube. 60 μL of $CuSO_4$ pentahydrate (7.4 mg/ml in water) was mixed with 60 μL BTTAA[4] (Click Chemistry Tools, 77.4 mg/ml in PBS) in a PCR tube, and 108 μL of this mixture was added to the reaction. After the mPEG dissolved completely, 1.5 mL GVs (OD 50) and 54 μL sodium ascorbate (59.4 mg/mL freshly made in PBS) were added. The mixture was rotated slowly overnight at room temperature and purified by 4 rounds of centrifugation (600xg, 2h). Other polymer formulations were similarly prepared.

#### GV characterization

Dynamic light scattering: A disposable cuvette containing 300 μL of a GV suspension diluted to approximately OD 0.2 was placed in a ZetaPALS particle analyzer and analyzed with the ZetaPALS Particle Sizing software using an angle of 90°, thin shell setting, run length of 15 s, and 6 runs per sample.

Zeta potential: An electrode (SZP, Brookhaven Instruments) was inserted into a mixture of 50 μL of OD 2 GVs and 1500 μL Milli-Q water in a disposable cuvette (BI-SCP, Brookhaven Instruments). Measurements were performed with a ZetaPALS particle analyzer (Brookhaven Instruments) running the ZetaPALS Zeta Potential software. Charge was calculated using the Smoluchowski model from 5 runs of 15 cycles.

Pressurized absorbance spectroscopy: 350μL OD 0.3 GVs were added to a flow-through quartz cuvette with a path length of 1 cm (Hellma Analytics). Hydrostatic pressure was applied from a 1.5 MPa nitrogen gas source through a single valve pressure controller (PC series, Alicat Scientific). Pressure was ramped from 0 to 900 kPa in 25 kPa increments with a 7 s equilibration period prior to measurement of absorbance at 600 nm with a microspectrometer (STS-VIS, Ocean Optics). Data were normalized to absorbance at 0 kPa and 900 kPa. Midpoint of collapse was derived from linear interpolation at normalized OD of 0.5.

TEM: GVs were exchanged into water and diluted to approximately OD 0.5. 3 μL of sample was spotted onto Formvar/carbon 200 mesh copper grids (Ted Pella) that were rendered hydrophilic by treatment with oxygen plasma (K100X, Emitech). After 90 s, sample was wicked off and stained for 1 min with 3 μL 1% uranyl acetate solution. Grids were imaged using a Tecnai T12 LaB6 120 kV transmission electron microscope (FEI Company) equipped with a Gatan UltraScan 2000 x 2000 CCD camera and DigitalMicrograph software interface (v3.9.0, Gatan Inc). Images were processed with FIJI[5].

MALDI-MS: Matrix-assisted laser desorption/ionization mass spectrometry (MALDI-MS) analysis of GV and GV-alkyne was performed with a Bruker Autoflex MALDI-time-of-flight mass spectrometer in linear positive ion mode. The GV sample was desalted by repeated buoyancy purification using 0.1% v/v trifluoroacetic acid (TFA) in water and concentrated to OD ~100. The GV sample was mixed 1:5 with sinapinic acid (10 mg/mL) dissolved in 30% acetonitrile + 0.01% TFA. The mixture was sonicated for 10 min and spotted onto the MALDI target plate.

**Small molecule and polymer synthesis**

Synthesis of CTA1 (**Fig. S13**): 4-Cyano-4-(ethylsulfanylthiocarbonyl) sulfanylvpentanoic acid N-hydroxysuccinimide ester[6] (600 mg, 1.66 mmol, 1 equiv) was dissolved in 10 mL dichloromethane, and 1-azido-3-aminopropane[7] (183 mg, 1.83 mmol, 1.1 equiv) and triethylamine (0.58 mL, 4.16 mmol, 2.5 equiv) were added. The mixture was stirred at room temperature for 14.5 h. The mixture was purified by column chromatography (3:2 hexanes:ethyl acetate) to yield the product as orange oil (404 mg, 1.17 mmol, 70.4% yield).

Synthesis of zwitterionic polymer (**Fig. S14**): The polymer was synthesized by reversible addition-fragmentation chain transfer (RAFT) polymerization. The monomer[8] (1.765 g, 5.010 mmol, 58 equiv), **CTA1** (30 mg, 0.086 mmol, 1 equiv), and AIBN (2.8 mg, 0.017 mmol, 0.2 equiv) were dissolved in DMSO (5.4 mL) and freeze-pump-thawed three times. The polymerization under argon atmosphere was initiated by immersing the flask into a 70 °C oil bath. The polymerization was quenched after 4.5 h by freezing with liquid nitrogen and exposing to air. The reaction mixture was precipitated into cold acetone, then the precipitate was redissolved in methanol and precipitated into cold ether. The precipitate was dried *in vacuo* to yield the *tert*-butyl ester protected polymer (926 mg). The carboxylate groups on the polymer were deprotected by stirring the polymer (750 mg) in 1.5 mL TFA at 23 °C for 1 h, then neutralized with NaOH solution.

The polymer was purified by dialysis in water (MWCO 3.5 kDa Spectra/Por regenerated cellulose membrane, Spectrum Chemical, New Brunswick NJ). The water was removed by lyophilization to yield the polymer.

Characterization: Mass spectrometry of **CTA1** was performed on a JEOL JMS-T2000GC AccuTOF GC-Alpha system using FD ionization. Infrared (IR) spectra were recorded on a Perkin Elmer Paragon 1000 spectrometer using neat samples on ATR diamond and are reported in frequency of absorption ($cm^{-1}$). $^{1}H$ NMR spectra were recorded on a Varian Inova 500 spectrometer (500 MHz), and $^{13}C$ NMR spectrum was recorded on a Bruker Ascend 400 spectrometer with Prodigy broadband cryoprobe (400 MHz)

**In vitro ultrasound imaging**

Images were acquired using a 128-element linear array probe (L22-14vX, Verasonics) with a center frequency of 18 MHz, elevation focus of 8 mm, elevation width of 1.6 mm, and element pitch of 0.10 mm. The transducer was connected to a programmable ultrasound scanner (Verasonics Vantage 128) operating on Vantage 4.4.0 software. Phantoms were cast out of 1% agarose in PBS using custom printed molds containing pairs of 2-mm diameter cylindrical wells. Samples for imaging were mixed 1:1 with 2% low melt agarose in PBS, incubated for 10 s at 42°C, and quickly loaded into the wells. After solidification, the phantoms were placed on top of an acoustically absorbent material and immersed in PBS to couple the sample with the imaging transducer. Samples were positioned such that the wells were at a depth of 8 mm.

Linear imaging was performed using a B-mode pulse sequence consisting of 89 ray lines, each transmitted from a 40-element aperture as a 2.5 cycle 18 MHz pulse with an 8-mm focus. Nonlinear imaging was performed with the same parameters using an amplitude modulation pulse sequence[9]. The imaging sequence for each sample consisted of 10 frames acquired at a transmit voltage of 1.6 V, 20 frames at 20 V, and 10 more frames at 1.6 V. Circular regions of interest were drawn in each well, and signal intensities were calculated by subtracting average pixel value across the last 10 frames from that of the first 10 frames.

**In vivo imaging**

All *in vivo* experiments were performed under protocols approved by the Institutional Animal Care and Use Committee at the California Institute of Technology. Imaging experiments were performed on 8-12 week old female BALB/cJ mice (Jackson Laboratory) and 12 week old male NSG mice (Jackson Laboratory).

Mice were maintained under 1.5% isoflurane anesthesia on a temperature-controlled imaging platform set to 38.5°C (Stoelting Co.) and head-fixed in a stereotaxic frame (Knopf). A PE10 catheter with a 30G needle was inserted into a lateral tail vein and the mouse was depilated with Nair, taking care to limit contact time to minimize scalp injury. A high-frequency transducer (L22-14vX, Verasonics) was coupled to the head through a column of ultrasound gel (centrifuged at 300xg, 20 min to remove bubbles) and positioned to capture a full coronal section at an arbitrary plane along the rostrocaudal axis.

Ultrafast power Doppler images were acquired at 15.625 MHz with a frame rate of 0.25 Hz as previously described[10,11]. Briefly, the pulse sequence consisted of 11 tilted plane waves varying from –10° to 10°. Each plane wave contained 4 cycles transmitted at a center frequency of 15.625 MHz with a voltage of 3 V. An ensemble of 250 coherently compounded frames, collected at a frame rate of 500 Hz, was processed through a singular value decomposition filter[12] (cutoff of 20) to produce a single power Doppler image. GVs (100 µL OD 50) were manually injected as a bolus 300 s after the start of acquisition. For multiple injection experiments, mice were given a second dose of GVs after 1 or 4 weeks.

Pixel-wise signal enhancement was calculated as the ratio of intensity at each time point relative to mean intensity in the first 75 frames. Time courses were extracted by averaging signal enhancement within a manually drawn region of interest encompassing the cortex. Each curve was smoothed using a 20-unit median filter. To calculate half-life, an exponential decay function was fitted to each curve from its maximum to the end of acquisition.

**RBC incubation**
Whole blood was collected from 8-12 week old female BALB/c mice by cardiac puncture using EDTA as an anti-coagulant. 500 µL of blood was aliquoted into 2 mL microcentrifuge tubes, diluted with 1.3 mL PBS, and centrifuged at 1000xg for 10 min. After discarding the supernatant, the cells were resuspended to approximately 1.8 mL in PBS and centrifuged again. After repeating this process once more, the RBCs were resuspended in PBS to approximately 4x the volume of the cell pellet (25% hematocrit).

Ultrasound imaging: 10 µL purified RBCs were mixed with 50 µL OD2 GVs which were urea-treated to produce nonlinear contrast. After incubating at room temperature for 1 h, samples were diluted with 900 µL PBS and centrifuged at 200xg for 8 minutes. The supernatant was removed, and this wash process was repeated once before concentrating the samples to approximately 60 µL. RBC only controls were prepared by mixing 10 µL RBCs with 50 µL PBS. GV only controls were prepared by mixing 50 µL OD2 GVs with 10 µL PBS. As described above, samples were loaded into phantoms and imaged using an amplitude modulation pulse sequence.

Flow cytometry: Purified RBCs were washed and resuspended to approximately 1e10 cells/mL in 1%w/v bovine serum albumin (BSA) in Hanks' Balanced Salt Solution (HBSS, Gibco). 10 µL OD10 GVs labeled with AF647, 10 µL RBCs, and 80 µL HBSS were combined in a 1.5 mL microcentrifuge tube. After incubating at 37°C for 1 h, samples were diluted with 1 mL HBSS and centrifuged at 600xg for 10 min. The supernatant was removed, and this wash process was repeated two more times before suspending in 100 µL 1% BSA in HBSS. RBCs were labeled with 0.2 µL FITC anti-mouse TER119 (BioLegend cat#116205) for 30 minutes at room temperature. Samples were washed by three rounds of centrifugation and resuspended in 1 mL 1%

BSA/HBSS. 50 µL of each sample was analyzed on a MACSQuant Analyzer 10 Flow Cytometer (Miltenyi Biotec) using the B1 and R1 channels for FITC-TER119 and AF647-GV, respectively. Data were analyzed in FlowJo. Single cells were isolated by FSC-H vs. FSC-A and displayed as R1-H vs. B1-H (see **Fig. S3**).

**Serum incubation**

To prepare sera from naïve BALB/c and NSG mice, blood was extracted by cardiac puncture, allowed to clot at room temperature for 15 min, centrifuged at 1000xg for 10 min, and carefully collected by pipette. GV antisera were similarly prepared from BALB/c mice 10 days after injection of 100 µL OD 50 Ana or Ana-PEG. Outbred mouse serum was purchased from Sigma Aldrich.

GVs (5 µL OD10) were mixed with 45 µL mouse serum and incubated undisturbed for 1 h at 37°C. Trans-illumination images were acquired before and after incubation using a Chemi-Doc gel imager (Bio-Rad). For ultrasound imaging, the samples were gently mixed, loaded into a phantom, and imaged as described above using a B-mode pulse sequence.

**LC-MS/MS**

GVs (100 µL OD50) were mixed with normal mouse serum (Sigma Aldrich) and incubated for 1 h at 37°C under gentle rotation. GVs were purified, exchanged into water, and concentrated over 4 rounds of centrifugation. 20 µL of these samples were transferred to 1.5 mL Low Protein Binding microcentrifuge tubes, resuspended in 980 µL of ultrapure water (Barnstead GenPure 18.2 $M\Omega cm^{-1}$), and centrifuged at 600xg for 30 min. After removing the subnatant, the GVs were resuspended to a total volume of approximately 74 µL with ultrapure water. Mouse serum controls were prepared by diluting 100 µL of the mouse serum into 900 µL of ultrapure water, then transferring 10 µL of this solution into three 1.5 mL Low Protein Binding microcentrifuge tubes (Sarstedt, Thermo Fisher) and diluting to 74 µL with ultrapure water. 10 µL of 500 mM ammonium bicarbonate, 10 µL of 10% (w/v) sodium deoxycholate (Fluka Analytical), and 4 µL of 250 mM dithiothreitol (Bio Basic) were added to each tube. The samples were incubated at 80°C for 10 minutes, then cooled to room temperature. 10 µL of 450 mM iodoacetamide was added to each sample. The samples were incubated for 30 minutes in the dark at room temperature, then 2 uL of Sequencing Grade Modified Trypsin (Promega) was added to each sample. After incubating overnight in the dark at room temperature, 100 µL of ethyl acetate followed by 5 µL of 10% (w/w) formic acid was added to each sample. The samples were vortexed for 1 minute to mix thoroughly, then centrifuged at 1000xg for 10 minutes to allow the aqueous and organic phases to fully separate. The organic (supernatant) phase was removed by pipette, and the aqueous layer was transferred to fresh 1.5 mL Low Protein Binding microcentrifuge tubes. Samples were stored at –20°C prior to characterization.

Digested peptides were submitted to the SPARC BioCentre (The Hospital for Sick Children, Peter Gilgan Centre for Research and Learning) for LC–MS/MS using an Orbitrap Fusion Lumos Tribrid Mass Spectrometer (Thermo Scientific). Data were analyzed using Proteome Discoverer v.2.5.0.400. Peptides were identified by searching against mouse (Uniprot_UP000000589_Mouse_15092020) UniProt reference databases.

A minimum protein confidence threshold of 95%, a minimum of 2 identified peptides, and a minimum peptide confidence threshold of 95% were used. Spectral counts and precursor intensity of each protein were analyzed using intensity-based absolute quantification (iBAQ) with Scaffold v.5.2.0 (Proteome Software Inc.). Gene Ontology Annotations for Mouse (https://www.ebi.ac.uk/GOA/downloads), downloaded October 10, 2022, were used to determine whether proteins were associated with immune response. Proteins annotated as part of an immunoglobulin complex, involved in complement activation, or involved in complement binding were deemed to be associated with immune response. Donut plots were created based on the geometric mean of abundance for each protein across the three replicates. The quantified protein data were log2-transformed and analyzed with Perseus v2.0.7.0 (MaxQuant).

A multiple unpaired t-test analysis using the false discovery rate method of Benjamini and Hochberg was used to identify differentially abundant proteins on the two gas vesicle types. Proteins identified with a false discovery rate of 5% and a log2 fold-change greater than 1 were deemed differentially abundant.

**Statistical analysis**
Sample sizes were chosen based on preliminary experiments to yield sufficient power for the proposed comparisons. Statistical methods are described in applicable figure captions.

**Data and code availability**
All gas vesicles, plasmids, data, and code are available from the authors upon reasonable request.

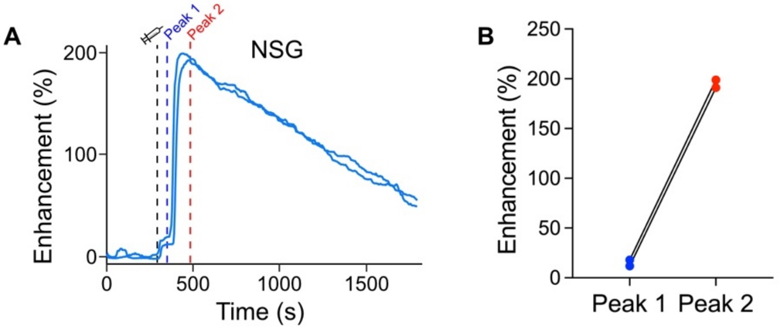


**Figure S1.** The second peak of hemodynamic contrast is independent of antibodies. **(A)** Time courses of Doppler signal enhancement following IV injection of 100 µL OD 50 Ana into NSG mice. N = 2. Dashed lines mark time of injection (black, 300 s), peak 1 (blue, 350 s), peak 2 (red, 480 s). **(B)** Enhancement at the indicated timepoints in time courses from panel A.

**A**

| | GV | RBC | Water |
|---|---|---|---|
| Radius, r (µm) | 0.062 | 2.215 | |
| Density, $\rho$ (kg/m$^3$) | 1.2 | 1110 | 1000, denoted as $\rho_w$ |
| Compressibility, $\kappa$ (m$^2$/N) | 7.65e-6 | 3.48e-10 | 4.5e-10, denoted as $\kappa_w$ |
| Wavenumber, $k$ (m$^{-1}$) | | | 10000 |
| Scattering cross section ($\sigma_S$, nm$^2$) of a single particle | 1.53e-7 | 1.83e-7 | |

$$\sigma_S = \frac{4\pi k^4 r^6}{9}\left[\left|\frac{\kappa-\kappa_w}{\kappa_w}\right|^2 + 3\left|\frac{\rho-\rho_w}{2\rho+\rho_w}\right|^2\right]$$

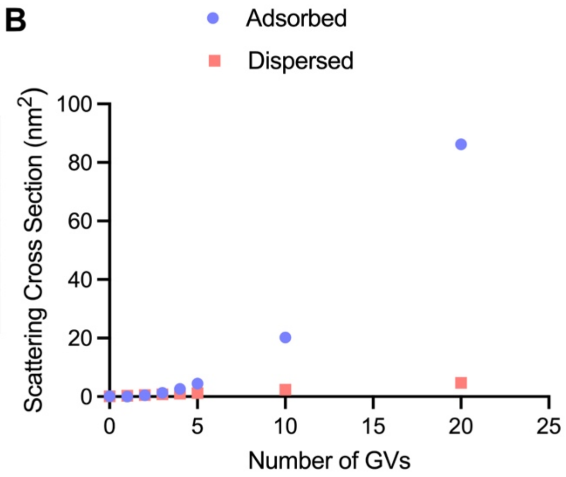


**Figure S2.** Modeling of scattering cross section. **(A)** Particles were modeled as uniform spheres using published volume[13], density[14], and compressibility[15,16] parameters. Density and compressibility of RBC-GV complexes were calculated using volume-weighted averages of the values for RBCs and GVs alone. The wavenumber was chosen based on a 15 MHz acoustic wave in a medium with a speed of sound of 1500 m/s. Scattering cross section of a Rayleigh scatterer was calculated using the equation shown[17]. **(B)** Scattering cross section of a single RBC with the indicated number of GVs. The dispersed case was calculated by summing the cross sections of individual particles, which assumes particles are dispersed enough to incoherently scatter. The adsorbed case was calculated based on a single particle with volume-weighted parameters.

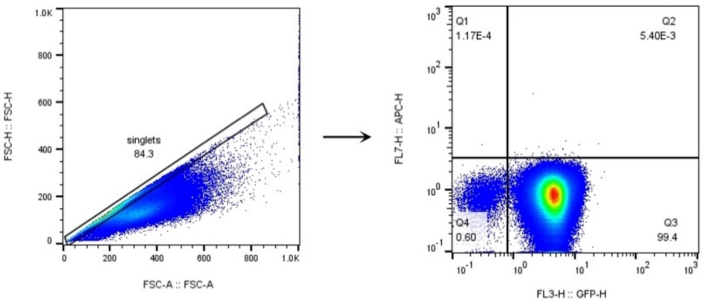


**Figure S3.** Gating strategy for flow cytometry of RBCs. Single cells were gated based on FSC-H vs FSC-A. Quadrants were defined based on an RBC-only control sample.

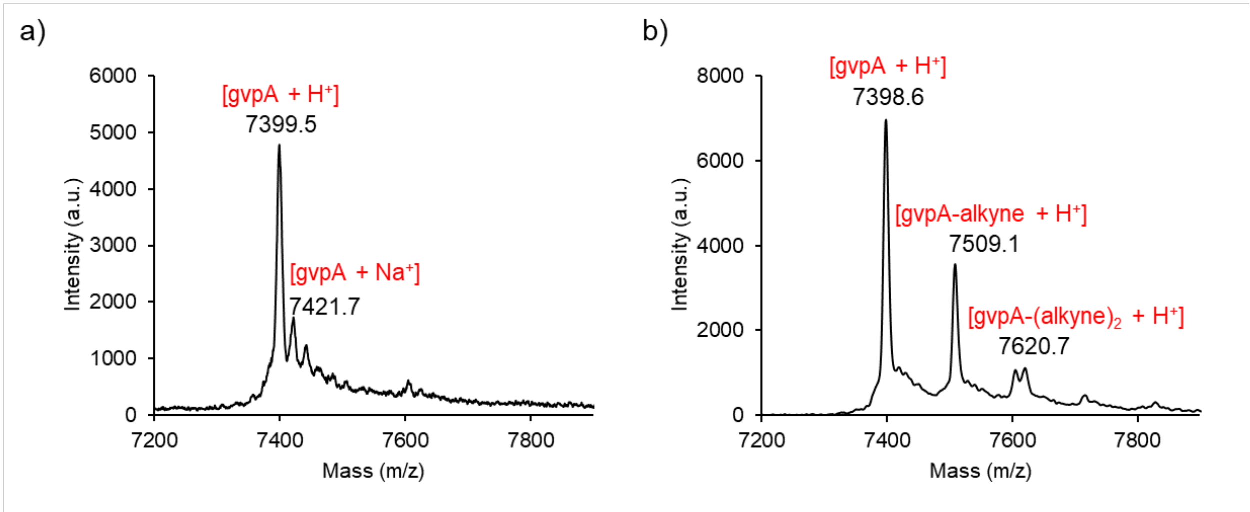


**Figure S4.** MALDI mass spectrum of Ana (**A**) and Ana-alkyne (**B**).

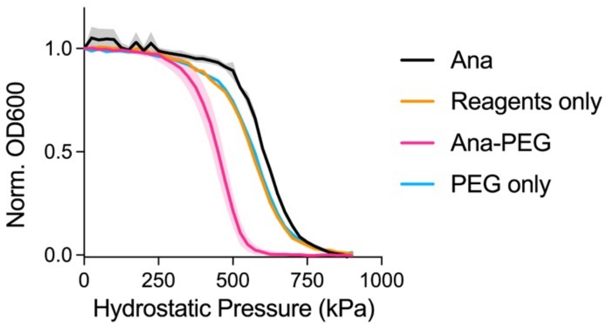


**Figure S5.** Hydrostatic collapse pressure of GVs mixed with CuAAC reagents in the absence of mPEG-azide or mixed with mPEG-azide without CuAAC reagents. N = 3. Curves for Ana and Ana-PEG shown for comparison. Thick lines, mean; shaded area, ± SEM.

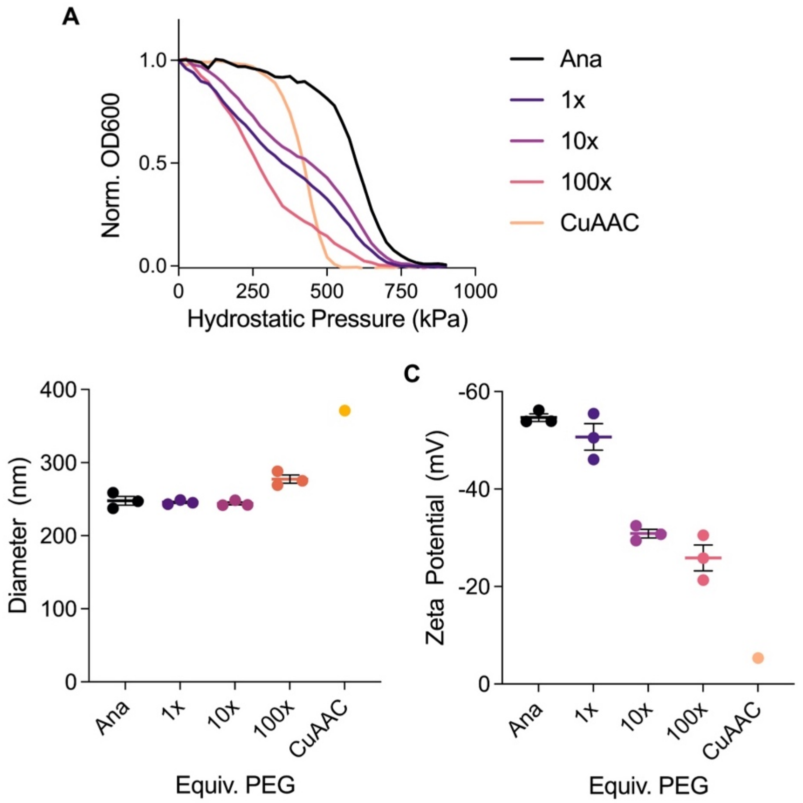


**Figure S6**. Direct functionalization with NHS-PEG. **(A)** Representative hydrostatic collapse pressure curves for GVs reacted with the indicated ratio of NHS-PEG, calculated relative to gvpA. Curves for unmodified Ana and Ana-PEG prepared by CuAAC are shown for comparison. N = 3. **(B)** Dynamic light scattering measurements, shown with Ana and Ana-PEG. N = 3. Error bars, ± SEM. **(C)** Zeta potential measurements, shown with Ana and Ana-PEG. N = 3. Error bars, ± SEM.

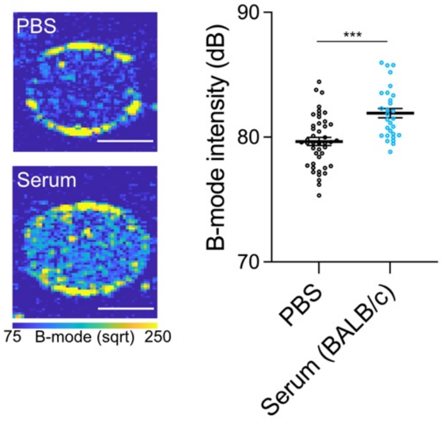


**Figure S7.** Ultrasound imaging of Ana GVs exposed to serum from naïve BALB/c mice. Left: Representative B-mode images after incubation at 37 °C for 1 h in PBS or serum. Right: Mean B-mode intensity within each well. N = 24. Error bars, ± SEM. Welch's t-test (***, $p < 0.001$).

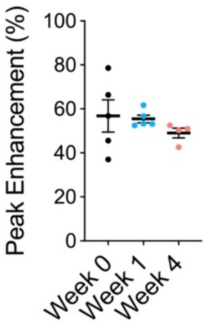


**Figure S8**. Peak signal enhancement in Ana-PEG time courses from Fig. 4E. Error bars, ± SEM.

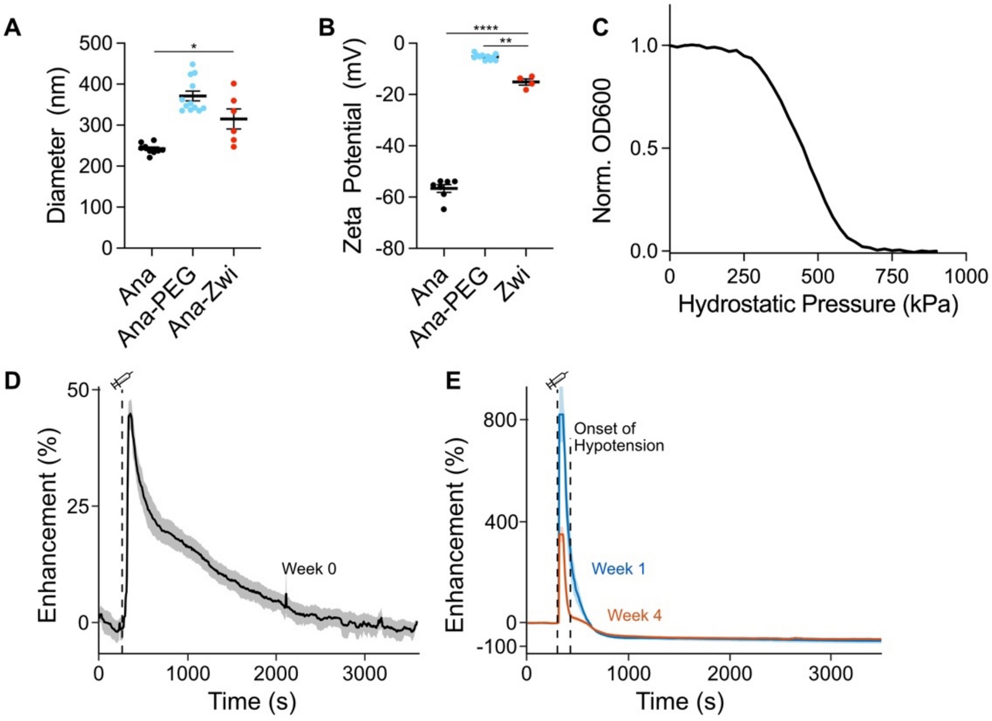


**Figure S9.** Functionalization with zwitterionic polymer does not reduce immune reaction. **(A)** DLS measurements of GVs coated with polyzwitterion. N = 6. Error bars, ± SEM. Welch's t-test, (*, $p < 0.05$). **(B)** Zeta potential measurements. N = 4. Error bars, ± SEM. Welch's t-test, (**, $p < 0.01$; ****, $p < 0.0001$). **(C)** Representative hydrostatic collapse pressure curve of Ana-Zwi. **(D)** Mean hemodynamic signal enhancement time course following IV injection of 100 µL 5.7 nM Ana-Zwi at 300 s (dashed line). N = 4. Shaded area represents ± SEM. **(E)** Mean hemodynamic signal enhancement following a second injection of Ana-Zwi after 1 (N = 2) or 4 weeks (N = 3). Dashed black line, time of injection (300 s); dashed red line, onset of hypotension (500 s); thick lines, mean; shaded area, ± SEM.

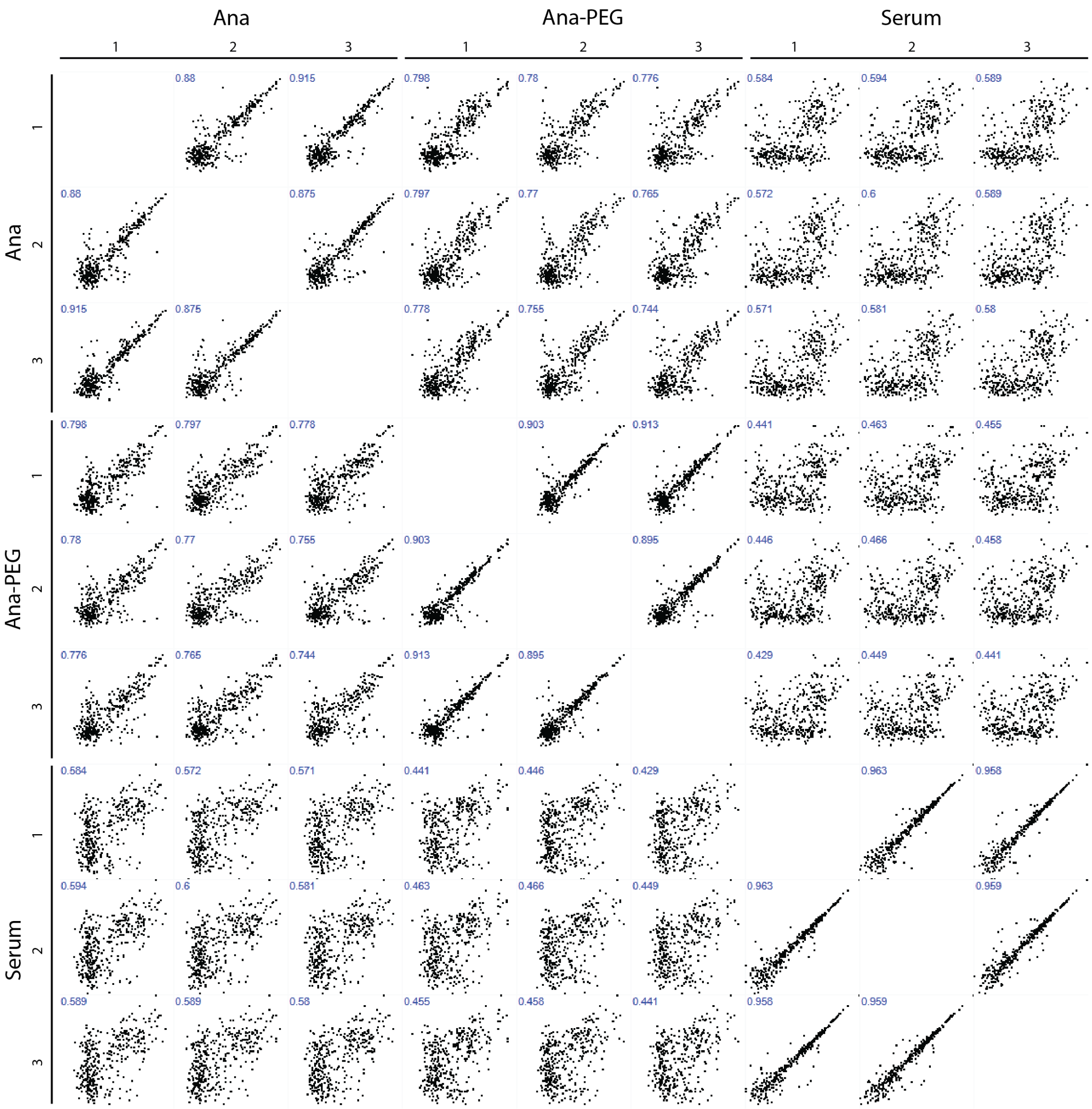


**Figure S10.** Multi-scatter plot of protein abundances in each sample. Pearson correlation coefficient (R) is shown in the top left of each subplot.

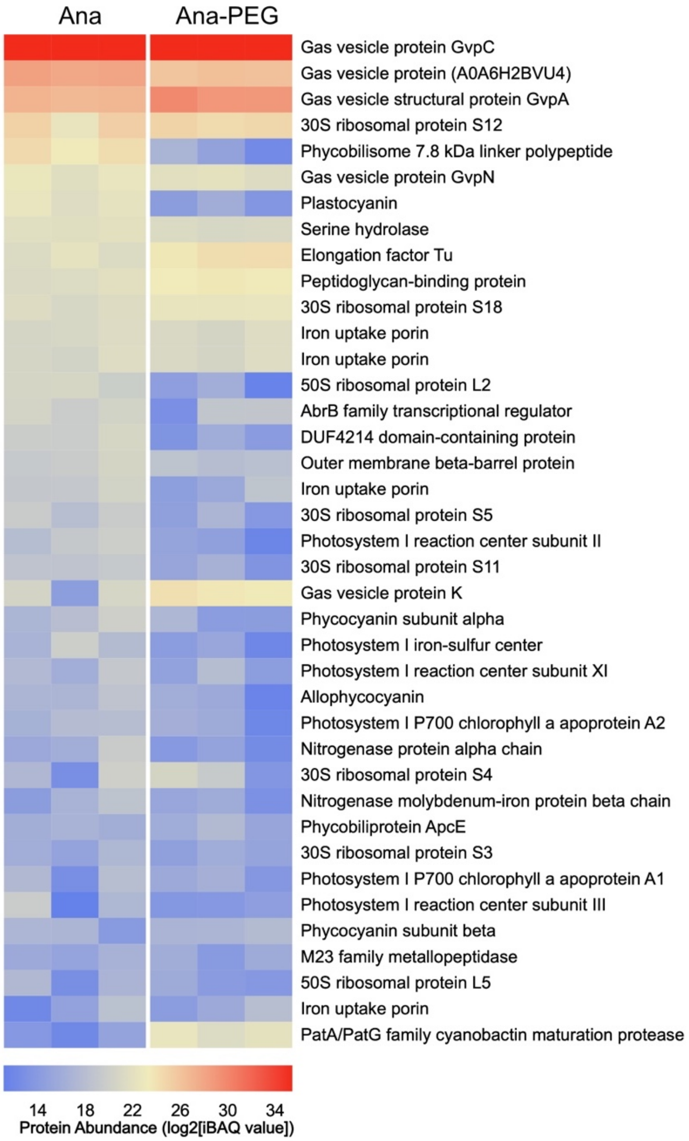


**Figure S11.** Heat map of cyanobacterial proteins detected by LC-MS/MS.

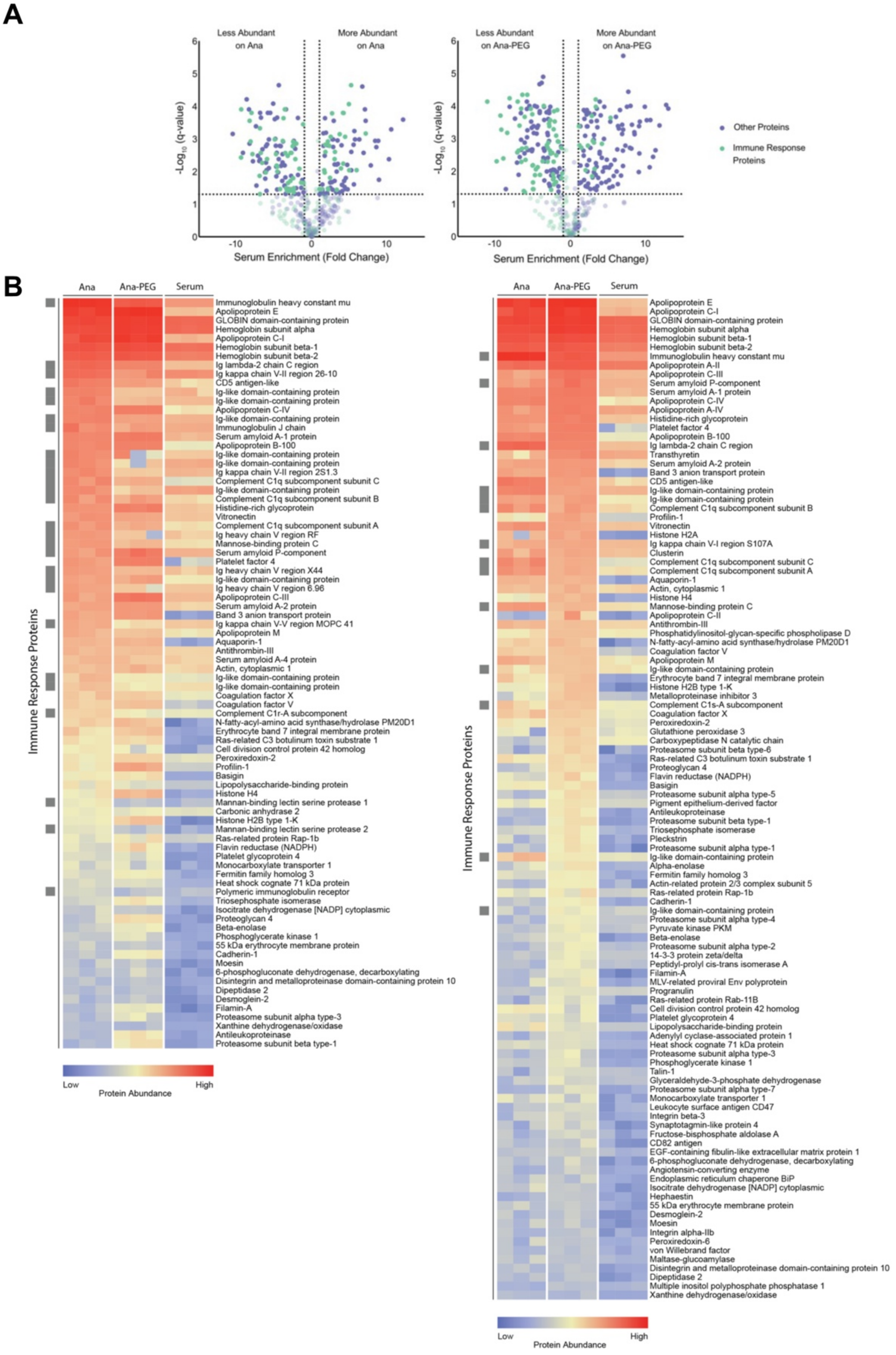


**Figure S12.** Protein abundances relative to serum**. (A)** Volcano plots of protein abundances on Ana (left) and Ana-PEG (right) compared to serum. Abundances were compared by multiple unpaired t-test analysis using the false discovery rate method of

Benjamini and Hochberg. Proteins with a false discovery rate of 5% and $log_2$ fold-change greater than 1 were deemed differentially abundant. Proteins that were not differentially abundant are shown translucently**. (B)** Heat map of differentially abundant proteins on Ana (left) and Ana-PEG (left) compared to serum. Immune response proteins are indicated by a gray box on the left.

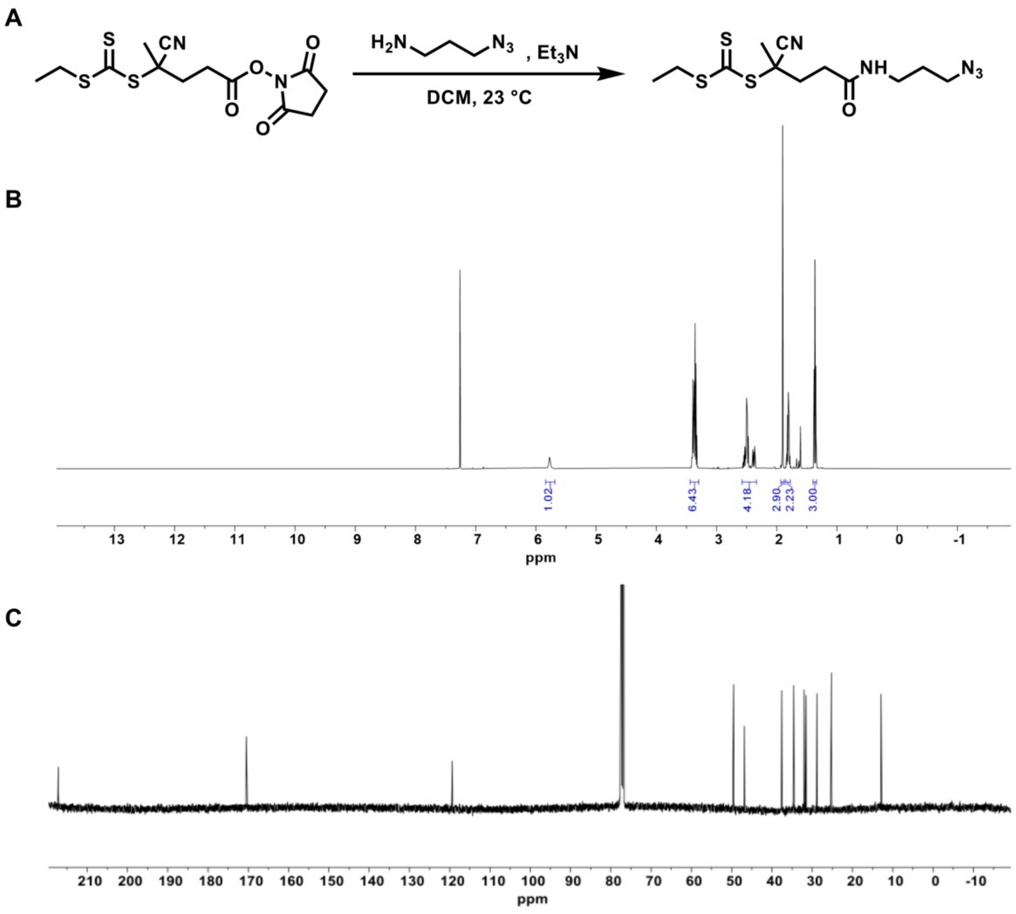


**Figure S13.** Synthesis of CTA1. **(A)** Reaction scheme. $^{1}H$ (**B**, 500 MHz) and $^{13}C$ (**C**, 101 MHz) NMR spectra of **CTA1** in $CDCl_3$. $^{1}H$ NMR (500 MHz in $CDCl_3$) δ: 5.77 (br s, 1H), 3.41–3.34 (m, 6H), 2.56–2.47 (m, 4H), 1.90 (s, 3H), 1.81 (p, *J* = 7.43 Hz, 2H), 1.37 (t, *J* = 6.60 Hz, 3H). $^{13}C$ NMR (101 MHz in $CDCl_3$) δ: 217.18, 170.51, 119.35, 49.53, 46.83, 37.58, 34.58, 32.01, 31.51, 28.85, 25.24, 12.90. MS $[C_{12}H_{19}N_5OS_3]^{+\bullet}$ calculated 345.07462, observed 345.07442. IR: 3292, 3084, 2929, 2871, 2093, 1645, 1548, 1447, 1375, 1260, 1152, 1116, 1073, 1031, 969, 942, 861, 801, 642 $cm^{-1}$.

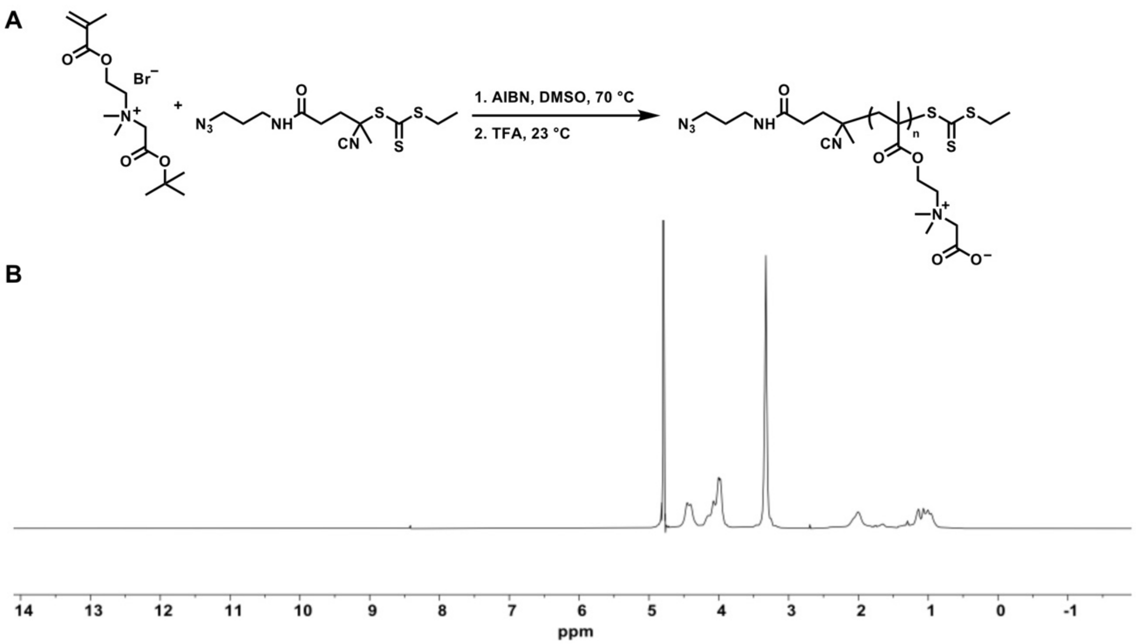


**Figure S14.** Synthesis of zwitterionic polymer. **(A)** Reaction scheme. (**B)** $^{1}$H NMR (500 MHz) spectrum of zwitterionic polymer in $D_2O$. $^{1}$H NMR (500 MHz in $D_2O$) δ: 4.55–4.30, 4.23–3.88, 3.45–3.21, 2.45–2.37, 2.17–1.88, 1.73–1.57, 1.46–0.83. $M_n$ = 16.3 kDa, $Đ$ = 1.25 (aqueous SEC). IR: 3382, 3027, 2966, 1722, 1623, 1472, 1452, 1421, 1386, 1335, 1263, 1234, 1147, 1058, 958, 936, 892, 845, 748, 712 cm$^{-1}$.

**Document S2**: LC-MS/MS dataset
